# Inhibition of Mitochondrial Fission Protein Drp1 Ameliorates Myopathy in the D2-mdx Model of Duchenne Muscular Dystrophy

**DOI:** 10.1101/2024.12.26.628172

**Authors:** H. Grace Rosen, Nicolas J. Berger, Shantel N. Hodge, Atsutaro Fujishiro, Jared Lourie, Vrusti Kapadia, Melissa A. Linden, Eunbin Jee, Jonghan Kim, Yuho Kim, Kai Zou

**Affiliations:** Department of Biology, University of Massachusetts Boston, Boston, MA; Department of Exercise and Health Sciences, University of Massachusetts Boston, Boston, MA; Department of Biomedical and Nutritional Sciences, University of Massachusetts Lowell, Lowell, MA; Department of Physical Therapy and Kinesiology, University of Massachusetts Lowell, Lowell, MA

**Keywords:** muscular dystrophy, mitochondria dynamics, Drp1, lipid peroxidation, muscle

## Abstract

Although current treatments for Duchenne Muscular Dystrophy (DMD) have proven to be effective in delaying myopathy, there remains a strong need to identify novel targets to develop additional therapies. Mitochondrial dysfunction is an early pathological feature of DMD. A fine balance of mitochondrial dynamics (fission and fusion) is crucial to maintain mitochondrial function and skeletal muscle health. Excessive activation of Dynamin-Related Protein 1 (Drp1)-mediated mitochondrial fission was reported in animal models of DMD. However, whether Drp1-mediated mitochondrial fission is a viable target for treating myopathy in DMD remains unknown. Here, we treated a D2-mdx model of DMD (9-10 weeks old) with Mdivi-1, a selective Drp1 inhibitor, every other day (i.p. injection) for 5 weeks. We demonstrated that Mdivi-1 effectively improved skeletal muscle strength and reduced serum creatine kinase concentration. Mdivi-1 treatment also effectively inhibited mitochondrial fission regulatory protein markers, Drp1(Ser616) phosphorylation and Fis1 in skeletal muscles from D2-mdx mice, which resulted in reduced content of damaged and fragmented mitochondria. Furthermore, Mdivi-1 treatment attenuated lipid peroxidation product, 4-HNE, in skeletal muscle from D2-mdx mice, which was inversely correlated with muscle grip strength. Finally, we revealed that Mdivi-1 treatment downregulated Alpha 1 Type I Collagen (Col1a1) protein expression, a marker of fibrosis, and Interleukin-6 (IL-6) mRNA expression, a marker of inflammation. In summary, these results demonstrate that inhibition of Drp1-mediated mitochondrial fission by Mdivi-1 is effective in improving muscle strength and alleviating muscle damage in D2-mdx mice. These improvements are associated with improved skeletal muscle mitochondrial integrity, leading to attenuated lipid peroxidation.

## INTRODUCTION

Duchenne muscular dystrophy (DMD) is an x-linked severe and progressive muscle wasting disorder that affects approximately 1 in 5,000 boys worldwide (48). DMD arises from a recessive mutation in dystrophin, a structural protein responsible for linking muscle cell membranes to the extracellular matrix (25), resulting in impaired myofiber membrane integrity that leads to muscle damage, degeneration and fibrosis. DMD patients develop muscle weakness and wasting at early ages (2-5 years old) (69), leading to severe respiratory and cardiac failure in early adulthood, and eventually premature death (1). Although current approved treatments (e.g., Elevidys, Duvyzat, and glucocorticoids) have proven to be effective in preserving muscle strength and function, they frequently come with serious side effects and/or has limited age range for treatment (55, 76). Therefore, there remains a strong need to identify novel therapeutic targets for developing additional therapies to treat DMD and improve quality of life in DMD patients.

Mitochondria play a vital role in energy homeostasis and muscle contraction by generating ATP (12). Dystrophin-deficiency in DMD renders the myofibers more susceptible to damage during muscle contraction, leading to excessive intramyocellular Ca^2+^ influx to mitochondria, which causes mitochondrial damage and dysfunction (50). Indeed, mitochondrial dysfunction is a well-known pathological hallmark of DMD and precedes muscle degeneration in DMD (33, 50, 57), suggesting mitochondrial dysfunction may play an early role in the development of myopathy in DMD. For example, impaired mitochondrial respiration and elevated Reactive Oxygen Species (ROS) emission were detected in skeletal muscle from D2-mdx mice as early as 4-week-old (33), which preceded skeletal muscle damage and necrosis (50). As such, mitochondria have emerged as a novel therapeutic target in the field of DMD research (13).

Mitochondria are dynamic organelles that undergo constant cycles of fusion and fission to adapt to the bioenergetic demands of their cellular environment (71). Balanced mitochondrial dynamics between fusion and fission is critical in maintaining mitochondrial quality and function (72). At the molecular level, mitochondrial fusion is primarily regulated by Optic atrophy 1 (OPA1), Mitofusion 1 and 2 (Mfn1 and Mfn2) (66). On the other hand, mitochondrial fission is primarily mediated by Dynamin-Related Protein 1 (Drp1), which is recruited from cytosol to mitochondria outer membrane upon activation by a group of specific adaptors, such as mitochondrial fission protein 1 (Fis1), mitochondrial fission factor (Mff), and mitochondrial dynamics proteins of 49 and 51 kDa (Mid49 and Mid51) (8, 66, 75). Although Drp1-mediated mitochondrial fission is essential in maintaining skeletal muscle function health (21, 24), overexpression of Drp1 caused impaired muscle growth (67), highlighting the importance of maintaining optimal level of Drp1-mediated mitochondrial fission in muscle growth. Emerging studies have shown that in various mouse models of Duchenne muscular dystrophy (DMD), skeletal muscle mitochondrial dynamics are disrupted at a young age, with a shift towards excessive mitochondrial fission and over-activation of Drp1 (30, 50, 51, 61, 62). The significance of Drp1-mediated mitochondrial fission in muscle degeneration in DMD was further supported by the evidence that loss of Drp1 reduced muscle degeneration and improved mobility in dystrophin-deficient worm and zebrafish models (26, 62). However, the therapeutic potential of targeting Drp1 in treating myopathy in DMD remains unclear.

Mitochondrial division inhibitor 1 (Mdivi-1) is a cell-permeable pharmacological inhibitor of Drp1-mediated mitochondrial fission, which prevents the recruitment of Drp1 to mitochondria (14). It is by far the most accessible and effective pharmacological inhibitor of Drp1. Importantly, the therapeutic potential of Mdivi-1 has been reported in various neurodegenerative disease models such as Amyotrophic Lateral Sclerosis and Alzheimer’s disease (46, 60). With regards to skeletal muscle, however, the evidence is scarce. Our recent work found that Mdivi-1 treatment improved mitochondrial fitness by rebalancing mitochondrial dynamics and attenuating cellular ROS content in skeletal muscle (40). In addition, Rexius-Hall et al., reported that treating myotubes with Mdivi-1 *in vitro* enhanced myofibril contractile production (59). Altogether, there is strong scientific evidence to support the idea that Mdivi-1 could be a viable approach to treat skeletal muscle myopathy in DMD.

In this study, we sought to examine the effects of Mdivi-1, a pharmacological inhibitor of Drp1-mediated mitochondrial fission, on mitochondrial quality, function and skeletal muscle health in D2-mdx mice. We hypothesized that D2-mdx mice would have imbalanced mitochondrial dynamics with elevated Drp1-mediated mitochondrial fission, reduced mitochondrial oxygen consumption, higher production of ROS and impaired skeletal muscle strength compared to the wildtype control mice. However, Mdivi-1 treatment would alleviate these defects in D2-mdx mice.

## MATERIALS AND METHODS

### Animal Care and Study Design

Male D2.B10-*Dmd*^mdx^/J and DBA/2J mice were purchased (The Jackson Laboratories, Bar Harbor, ME. Stock ID #0013141 and #000671) at 4 to 5-weeks of age and acclimatized to the animal facility for 1 week. Animals were housed in a temperature and humidity-controlled environment and maintained on a 12:12 h light–dark cycle with food and water provided ad libitum.

After being acclimatized, D2.B10-*Dmd*^mdx^/J (D2-mdx) were randomly divided into either a vehicle (VEH, 2% DMSO in PBS) or Mdivi-1 treatment group (40mg/kg body weight Mdivi-1). DBA/2J (wildtype, WT) mice also received vehicle injections and served as the control group. Animals received intraperitoneal injections 3 times per week for 5 weeks (Figure 1). These interventions created 3 experimental groups: WT (n=8), D2-mdx/VEH (n=8), and D2-mdx/Mdivi-1 (n=8). This dose of Mdivi-1 has been previously reported to be safe in mice up to 8 weeks of treatment (3, 4). Mice were subjected to muscle function testing before and after the intervention to determine muscle strength. All Mice were euthanized 24 hours after the last injection.

**Figure 1:**
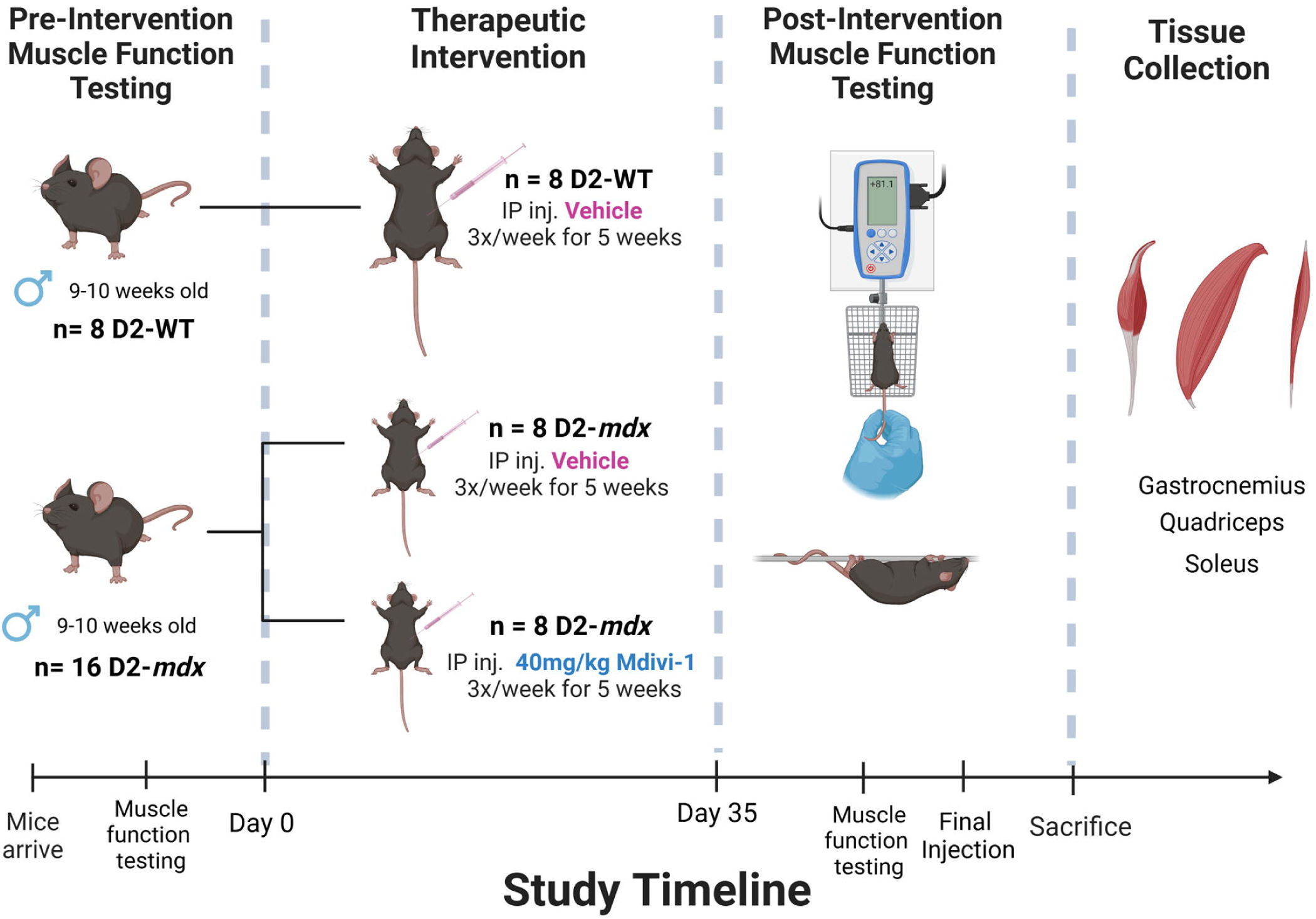
Study design schematic. (Illustrated using BioRender)

In addition, to evaluate whether Mdivi-1 had any adverse effects on phenotypes in normal healthy mice, we added another group with Mdivi-1 injections in wildtype mice (WT/Mdivi-1, n=4) and measured all functional tests. Due to the fact that Mdivi-1 treatment did not result in any detrimental effects on body phenotype and muscular function, we excluded WT/Mdivi-1 group from the rest of the mitochondrial and biochemical analyses and focused on the therapeutic effects of Mdivi-1 on D2-mdx mice as a treatment.

All experimental procedures were approved by the Institutional Animal Care and Use Committee of the University of Massachusetts Boston.

### Mdivi-1 Preparation

Mdivi-1 was purchased from Caymen Chemical (Ann Arbor, Michigan) and 100mg/mL stock solution was made using 100% DMSO. For injections, Mdivi-1 was diluted in sterile PBS (2mg/ml). Due to the poor aqueous solubility of Mdivi-1, each dose was gently sonicated in order to produce a homogenous suspension and delivered immediately through intraperitoneal injection as previously described (40, 58)

### Grip Strength Testing

The day before the last injection, mice were subjected to a grip strength test to determine limb muscle strength. All mice were placed on the wire grid of the BIOSEB BIO-GS4 Grip Strength Test meter (Bioseb, Pinellas Park, FL). Once all four limbs were gripping the grid, the mouse was gently pulled by the base of the tail and the peak pull force (g) was recorded on the digital force transducer. The peak pull force was collected for each mouse for 3 trials, with a 60 second rest period in between each trial. The output was recorded as force (g)/body weight (g). The average of the three trials was calculated.

### Hang Wire Testing

All mice were subjected to a hang wire test to determine limb strength and endurance as previously described (15). All mice were gently placed on the wire set up 12 inches from the base of the cage. Mice were left suspended on the wire until they reached exhaustion and dropped to the base of the cage. The time they remained suspended was recorded for three trials. All mice were given 60 second rest times between each trial. Impulse (s*g) was calculated according to DMD_M.2.1.004 standard operating procedures by multiplying the average time suspended (in seconds) by body mass (in grams).

### Tissue Collection

24 hours after the final Mdivi-1 injection, mice were euthanized using CO_2_ asphyxiation/cervical dislocation. Blood was collected immediately via cardiac stick and centrifuged for 15 minutes at 3,000 rpm at 4°C to collect serum. Quadriceps, soleus, gastrocnemius, and tibialis anterior muscles were collected, weighed, and stored for further analyses.

### Serum Creatine Kinase Activity

Serum creatine kinase activity was determined using a commercially available assay kit (ab155901, Abcam, Waltham, MA) and Biotek Synergy H1 Microplate Reader (Agilent, Lexington, MA). The protocol was completed per manufacturer’s instructions.

### Skeletal Muscle Mitochondrial Isolation

The quadricep was dissected from the mouse and was immediately added to 1 mL ice cold Mitochondrial Isolation Buffer 1 or IBM1 (67mM sucrose, 50mM Tris/HCl, 50mM EDTA/Tris, and 0.2% BSA) in a 5mL Eppendorf tube. Dissection scissors were used to snip muscle tissue until desired consistency was achieved. Sample was then transferred to 15mL conical tube and final volume was brought up to 5mL and 2.5uL trypsin was added (0.05% trypsin). Sample was incubated in trypsin for 45 minutes. After digestion, the sample was centrifuged at 200g, 4°C, for 3 minutes. After spin, supernatant was discarded, and pellet was resuspended in 3mL of IBM1 and then transferred to a 10mL Teflon glass homogenization tube. Tissue was homogenized using a drill press with serrated tissue grinding pestle attached (510rpm with 10-14 passages). After homogenization, homogenate was transferred to 15mL conical tube and total volume was brought up to 8mL using ice cold IBM1. The homogenate was centrifuged at 700g for 10 minutes at 4°C. Supernatant was transferred to a 38.5 ultra-clear tube and was centrifuged at 10,000g for 10 minutes at 4°C. Supernatant was discarded and pellet was resuspended in 150uL or ice-cold Mitochondrial Isolation Buffer 2 or IBM2 (250mM sucrose, 3mM EGTA/Tris, 10mM Tris/HCl). The sample was centrifuged again at 10,000g for 10 minutes at 4°C and IBM2 was used to resuspend mitochondria. After mitochondria isolation, protein concentration was determined using a Pierce BCA protein assay kit (Thermo Fisher Scientific, Waltham, MA).

### Mitochondrial Respiration

Isolated mitochondria were used to determine mitochondrial respiration rates by measuring oxygen consumption rates (OCR) with Seahorse XFp Extracellular Flux Analyzer (Agilent Technologies, Santa Clara, CA) as previously described (42). Immediately after protein quantification, isolated mitochondria were plated on the Seahorse plate at a concentration of 4 μg/well in the presence of 10 mM pyruvate and 5 mM malate. ADP (5 mM), oligomycin (2 μM), carbonyl cyanide-4 phenyl-hydrazone (FCCP, 4 μM), and antimycin (4 μM) were subsequently injected into ports to measure OCR under different respiratory states: Pyruvate+Malate to measure state 2 respiration rate, ADP (5 mM) to measure state 3 respiration rate, oligomycin (2 μM) to measure state 4 respiration rate, carbonyl cyanide-4 phenylhydrazone (FCCP, 4 μM) to measure maximal respiration rate, and antimycin (4 μM) to measure non-mitochondrial respiration rates. Respiratory control ratio (RCR) was calculated by state 3 respiration rate ÷ state 4 respiration rate and used to assess mitochondrial integrity. RCR is a measure used to assess efficiency of mitochondrial respiration and is calculated by dividing the rate of oxygen consumption with ADP stimulated respiration (state 3) by the respiration after oligomycin addition (state 4). Coupling efficiency is the proportion of oxygen consumed to drive ATP synthesis compared with that driving proton leak and is calculated as: (basal respiration-state 4 respiration)/basal respiration. Spare capacity is calculated as the difference between the maximal respiration and the basal respiration. All data were analyzed using the Agilent Seahorse Wave software.

### Mitochondrial Hydrogen Peroxide Production

Mitochondrial-derived H_2_O_2_ production (*m*H_2_O_2_) was measured fluorometrically as previously described (42). Briefly, *m*H_2_O_2_ was measured in Buffer Z (105 mM K-MES, 30 mM KCl, 1 mM EGTA, 10 mM K_2_HPO_4_, 5 mM MgCl_2_-6H_2_O, 2.5 mg/mL BSA, pH 7.1), supplemented with creatine (5 mM), creatine kinase (20 U/mL), phosphocreatine (30 mM, to mimic resting condition), Amplex Ultra Red (10 µM), horseradish peroxidase (20 U/mL), superoxide dismutase (20 U/mL), ATP (5 mM), and auranofin (0.1 µM). The following substrates assessed various sites: (1) pyruvate (10 mM) + malate (5 mM) to assess Complex I via generation of NADH; (2) pyruvate (10 mM) + malate (5 mM) + antimycin (2 µM) for the assessment of Complex III; (3) succinate (10 mM) + rotenone (4 µM) to assess Complex II via generation of FADH and (4) pyruvate (5mM) + rotenone (4µM) to assess pyruvate dehydrogenase complex (PDC) (56). All reactions were done at 37 °C, in a microplate reader (Thermo Fisher Scientific, Waltham, MA). Fluorescence values were converted to picomoles of H_2_O_2_ via an H_2_O_2_ standard curve, and H_2_O_2_ emission rates were calculated as picomoles of H_2_O_2_ per minute per milligram mitochondria (73).

### Transmission Electron Microscopy

Fresh skeletal muscle tissue (soleus) was immediately fixed in 2.5% glutaraldehyde in 0.1 M Sodium Cacodylate buffer (pH 7.2) for 24 hours at 4 °C and postfixed in 1% osmium for 1 h. Fixed tissues were dehydrated in a series of ascending ethanol concentrations, followed by two propylene oxide baths, and infiltrated using resin SPI-Pon 812 resin mixture per instructions and then switched to Resin/100% Propylene Oxide mixture (1:1), to polymerize overnight at 60 °C. Thin sections (70 nm) of polymerized Epon–Araldite blocks were cut using a Leica Ultracut UCT ultramicrotome placed on Cu grids (200 mesh size), and stained for 5 min in uranyl acetate, followed by 2 min in lead citrate. Muscle fibers were examined on a FEI (Thermo Fisher Scientific, Waltham, MA) Tecnai Spirit 12 transmission electron microscope and images captured using a Gatan Rio9, 9-megapixel side-mounted digital camera. Ten representative micrographs from subsarcolemmal and intermyofibrillar regions were acquired at ×19,000 magnification. Quantification was achieved using the ImageJ software.

### Mitochondrial Morphology Analysis

Mitochondrial morphology analysis was completed using a previously developed protocol by Lam et al. (43). Briefly, damaged mitochondria were determined by identifying mitochondria with visible damage, represented in TEM images as mitochondria with areas of white space. The ratio of damage is expressed as # of damaged mitochondria/ total # of mitochondria counted in image. Circumference, area, roundness, and aspect ratio parameters were calculated by ImageJ by tracing along the membrane of each individual mitochondria in each TEM image. Aspect ratio refers to the ratio of the length of a mitochondrion to its width, indicating how elongated each mitochondrion is; a lower aspect ratio indicates a more rounded or punctate mitochondrion, suggesting mitochondrial fragmentation.

### Immunoblot Analyses

Gastrocnemius muscles were homogenized in ice-cold homogenization buffer supplemented with protease and phosphatase inhibitors as previously described (42). Protein concentration was determined by using a Pierce BCA protein assay kit (Thermo Fisher Scientific, Waltham, MA). Equal amount of protein was loaded to Midi Protean Precast 4-15% gradient TGX Gels (Bio-Rad, Portland, ME), subject to SDS-Page, and transferred onto Nitrocellulose Membranes (Bio-Rad, Portland, ME) using Trans-Blot Turbo Transfer System (Bio-Rad, Portland, ME). Membranes were blocked in 5% BSA in 0.1% TBST and probed with a primary antibody (See the full list of antibodies in Supplemental Table 1) overnight at 4°C. Membranes were then probed by Horseradish peroxidase (HRP)-linked secondary antibodies (Cell Signaling, Danvers, MA) for one hour and developed under SignalFire ECL Reagent (Cell Signaling, Danvers, MA) solution before imaging with FluorChem M System imager. All images were quantified using ImageJ. Data were normalized to total protein content using Ponceau S staining.

### Quantitative real-time PCR (RT-PCR)

RNA was extracted using RNeasy kit (Qiagen, Hilden, Germany) as previously described (41). Concentrations and purity of RNA samples were assessed on a Biotek Synergy H1 Microplate Reader (Agilent, Lexington, MA). cDNA was reversed transcribed from a 100ng of RNA using a High-Capacity cDNA Reverse Transcription Kits (Applied Biosystems, Foster City, CA) following manufacture instructions. cDNA was amplified in a 0.2 mL reaction containing SYBR Green PCR Master Mix (for GPx4) or TaqMan Universal PCR Master Mix, TaqMan Gene Expression Assay and RNase-free water. RT-PCR was performed using a QuantStudio 3 Real-Time PCR (Thermo Fisher Scientific, Waltham, MA), and results were analyzed using Design and Analysis Application (Thermo Fisher Scientific, Waltham, MA). Gene expression was quantified for all genes of interest (Supplementary Table 2 and 3) using the ΔΔCT method. The expression of β-Actin (for GPx4) or GAPDH was used as the housekeeping gene.

### Statistical Analysis

Data were expressed as Mean ± SEM. For muscle functional data, Drp1 and Fis1 protein data, Two-way ANOVA was performed (main effect of *mdx and Mdivi-1*), followed by Fisher’s LSD post-hoc analysis when significant interaction was detected. For other data, One-way ANOVA was performed, followed by Fisher’s LSD post-hoc analysis when significant main effect was detected. Pearson correlation analysis was used to assess linear relationships between 4-HNE and grip strength, hangwire time or Col1a1 protein expression. All statistical analysis was performed using GraphPad Prism 10 with a significance level set at as P < 0.05.

## RESULTS

### Mdivi-1 treatment improves muscular strength and attenuates muscle damage in D2-mdx mice

We first sought to verify the phenotypical atrophy and muscle weakness reported in D2- mdx mice. When comparing with their respective WT controls, D2-mdx mice had significantly lower body weight throughout the course of study (Fig. 2A, main effect of mdx, P <0.0001) and lower weights of gastrocnemius, tibialis anterior, quadriceps, and heart (Fig. 2B, main effect of mdx, P <0.0001, P = 0.0008, P <0.0001, and P = 0.0002, respectively). However, Mdivi-1 treatment had no significant effect on body weight or any of these muscle tissue weights (Fig. 2A and B).

**Figure 2:**
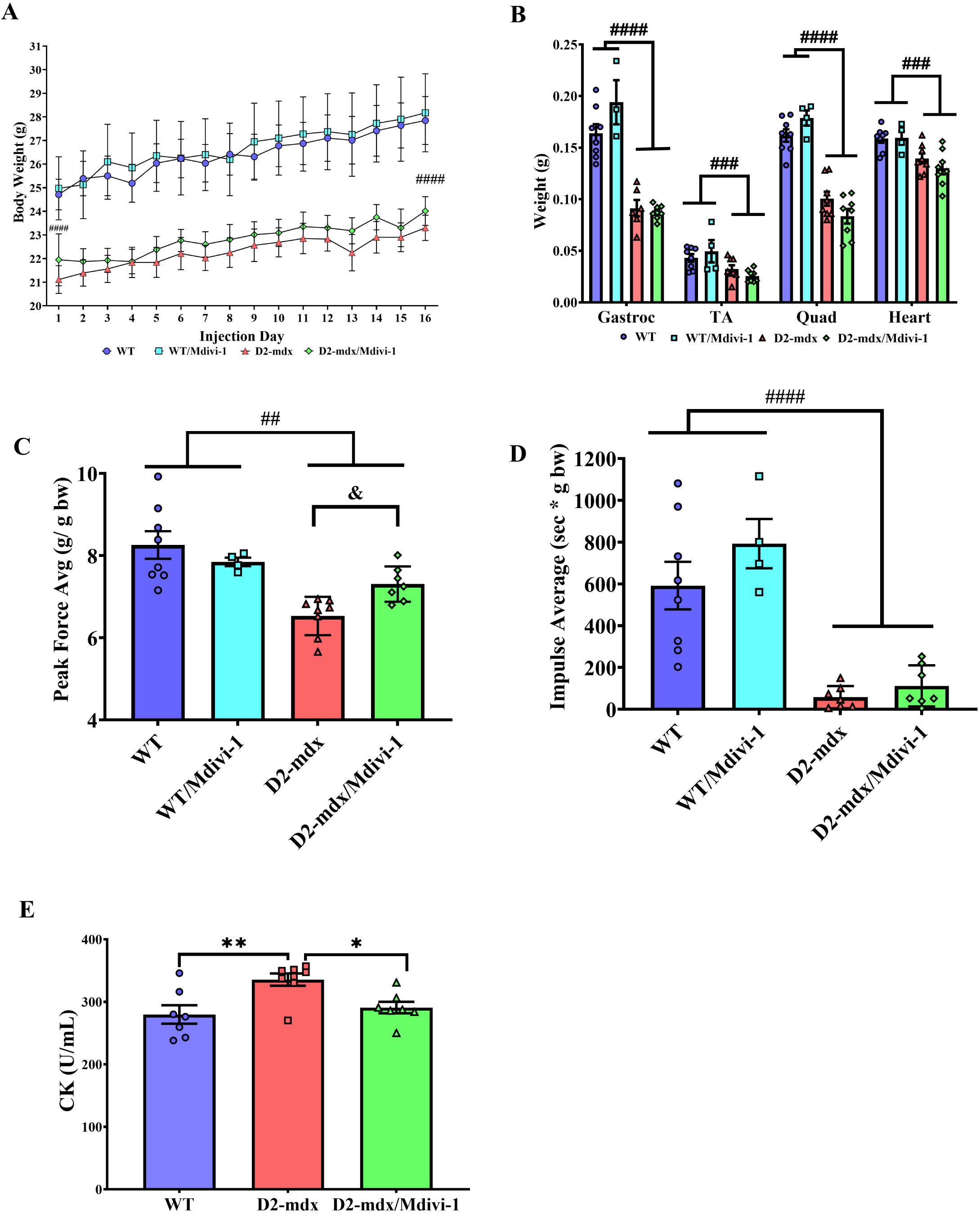
Mdivi-1 treatment improves muscular strength and attenuates muscle damage in D2-mdx mice. A) Body weights over a 5-week period. B) Tissue weights were reported at the time of tissue collection expressed as absolute weight (grams). Gastrocnemius (Gastroc), tibialis anterior (TA), quadriceps (Quad), and heart; C) Grip strength expressed in average peak force/body weight; D) Hang wire impulse testing expressed in time (sec) x body weight (grams). E) Creatine kinase activity assay from serum collected via cardiac stick. Data are presented as Mean ± SEM. N=4-8 mice per group. * P < 0.05; ** P < 0.01 significant difference between groups. ## P < 0.01; ### P < 0.001 and #### P < 0.0001 significant main effect of DMD. & P < 0.05 significant main effect of Mdivi-1.

Furthermore, D2-mdx mice had a lower grip strength when compared to WT controls (Fig. 2C, main effect of mdx, P = 0.010). Importantly, a Mdivi-1 and genotype interaction was noted in grip strength, revealing a significant improvement in grip strength in D2-mdx mice (14.3%, Fig. 2C, P = 0.046), but not in WT mice. Similarly, holding impulse (hang wire time normalized to body weight) also reflected a significant reduction in D2-mdx compared to WT mice (Fig. 2D, P <0.0001), and although there was not a statistically significant effect of Mdivi-1 treatment, a 92.2% improvement in holding impulse in D2-mdx/Mdivi-1group was found when compared to D2-mdx group (Fig. 2D). Finally, serum creatine kinase (CK) activity, a marker of muscle damage, was higher in D2-mdx mice compared to WT (Fig 2E, P = 0.006), but was significantly reduced in D2-mdx mice treated with Mdivi-1 when compared to the vehicle treated counterparts (Fig. 2E, P = 0.043).

### Mdivi-1 inhibits Drp1-Mediated mitochondrial fission machinery in skeletal muscle from D2-mdx mice

We next examined the effects of Mdivi-1 on protein markers of mitochondrial dynamics in skeletal muscle. We first confirmed that Mdivi-1 treatment inhibited Drp1 activation and total Drp1 content in skeletal muscle. Both Drp1(Ser616) phosphorylation and total Drp1 content were higher in skeletal muscle of D2-mdx mice (Fig. 3A and B, main effect of mdx, P = 0.003 and 0.049, respectively) when compared to WT, which were significantly attenuated with Mdivi- 1 treatment regardless of disease status (Fig. 3A and B, main effect of Mdivi-1, P = 0.003 and 0.005). Consistently, when normalized to total Drp1 content, there remained a significant reduction in the ratio of pDrp1 (Ser616)/Total Drp1 with Mdivi-1 treatment in both WT and D2- mdx mice (Fig. 3C, main effect of Mdivi-1, P=0.041). We next examined the effects of Mdivi-1 on the expression of several mitochondrial fission adaptor proteins in skeletal muscle. Consistent with Drp1 phosphorylation result, Fis1 was also significantly elevated in skeletal muscle of D2- mdx mice compared to WT mice (Fig. 3D, P < 0.0001), but was attenuated with Mdivi-1 treatment (Fig. 1C, P = 0.029). In addition, D2-mdx mice treated with Mdivi-1 had lower expression of Mitochondrial Fission Process 1 (MTFP1) in skeletal muscle compared to WT controls (Fig. 3D, P = 0.006), but not D2-mdx mice. With regards to mitochondrial fusion, D2- mdx mice had higher protein expression of Mfn1, but not Mfn2 or OPA1, in skeletal muscle compared to WT mice (Fig. 3D, P = 0.001). Interestingly, Mdivi-1 treatment reduced Mfn2 protein expression in D2-mdx mice (Fig. 3E, P = 0.042).

**Figure 3:**
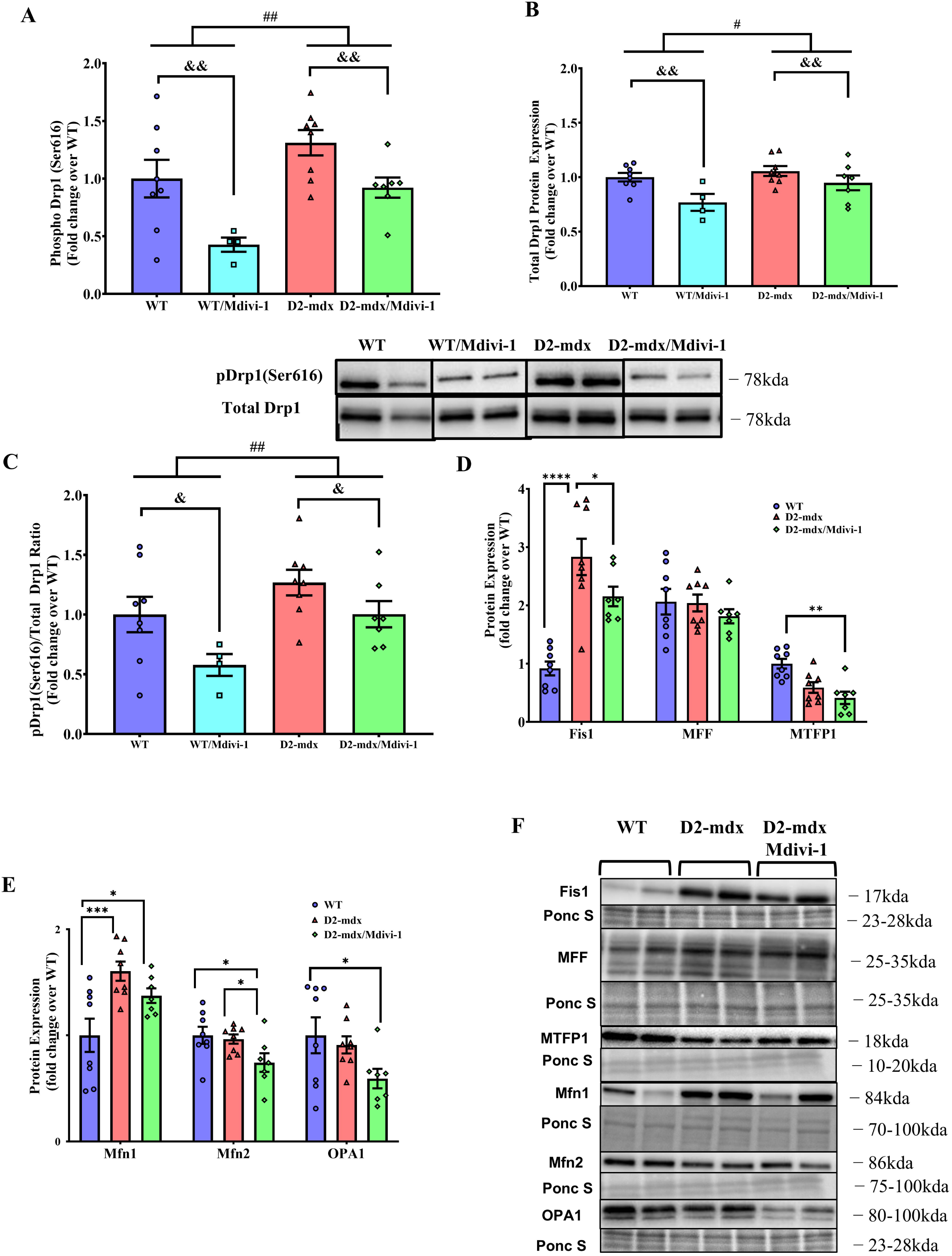
Mdivi-1 inhibits Drp1-Mediated mitochondrial fission and improves mitochondrial dynamics in skeletal muscle from D2-mdx mice. A) Drp1 (Ser616) phosphorylation. B) Total Drp1 protein expression. C) Ratio of phospho Drp1 (Ser616) over total Drp1. D) Protein expression of mitochondrial fission markers. E) Protein expression of mitochondrial fusion markers. F) Representative immunoblots for D and E. Data are presented as Mean ± SEM. N=4-8 mice per group. * P < 0.05; ** P < 0.01; *** P < 0.001; **** P < 0.0001 significant difference between groups. # P < 0.05; ## P < 0.01 significant main effect of DMD. & P < 0.05; & P < 0.01 significant main effect of Mdivi-1.

### Mdivi-1 treatment lowers autophagy markers, but not mitophagy, mitochondrial biogenesis or content markers in skeletal muscle from D2-mdx mice

D2-mdx mice had higher LC3B I and II protein content, but lower LC3BII/I ratio in skeletal muscle when compared to WT mice (Fig. 4A, P = 0.031, 0.028 and 0.005 respectively). Importantly, Mdivi-1 treatment significantly reduced LC3B I and elevated LC3B II protein content, which resulted in a restored LC3B II/I ratio in skeletal muscle from D2-mdx mice with no difference comparing to WT mice (Fig. 4A, P = 0.037). These results suggest that Mdivi-1 treatment was able to restore the autophagosome formation process in D2-mdx mice.

**Figure 4:**
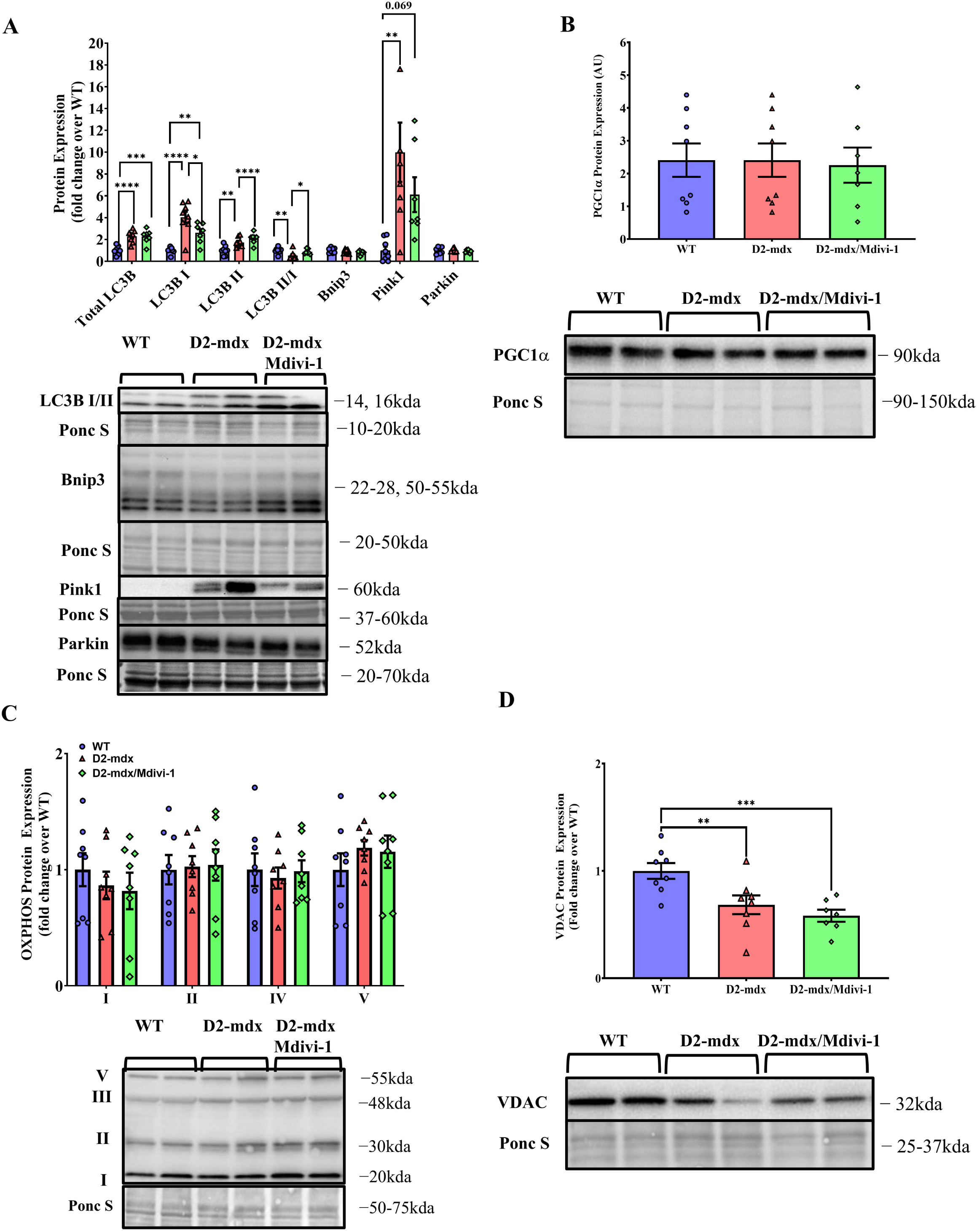
Mdivi-1 treatment lowers autophagy markers, but does not alter mitophagy, mitochondrial biogenesis or content markers in skeletal muscle from D2-mdx mice. A) Protein expression of autophagy and mitophagy markers. B) PGC1α protein expression. C) Protein expression of oxidative phosphorylation complexes. D) VDAC protein expression. Data are presented as Mean ± SEM. N= 7-8 mice per group. * P < 0.05; ** P < 0.01; *** P < 0.001; **** P < 0.0001 significant difference between groups.

Meanwhile, Pink1, a mitophagy regulatory marker that normally accumulates on the membranes of damaged mitochondria (2), was markedly higher in skeletal muscle from D2-mdx mice compared to WT controls (Fig. 4A, P = 0.002). However, there was no significant reduction of Pink1 with Mdivi-1 treatment. In addition, there were also no differences in protein expression of other mitophagy markers (i.e., Parkin and Bnip3), mitochondrial biogenesis marker PGC111, or oxidative phosphorylation complexes (OXPHOS) between groups (Fig. 4B-C). Lower protein expression of VDAC was exhibited in skeletal muscle in D2-mdx mice (Fig. 4D, P = 0.067) and remained lower with Mdivi-1 treatment (Fig. 4D, P = 0.001).

### Mdivi-1 treatment improves subsarcolemmal, but not intermyofibrillar mitochondrial morphology in skeletal muscle from D2-mdx mice

We next sought to assess whether the alterations in mitochondrial quality control regulatory machinery by Mdivi-1 treatment led to improvement in skeletal muscle mitochondrial morphology. The percentage of damaged subsarcolemmal mitochondria was higher in skeletal muscle from D2-mdx mice compared to WT (Fig. 5A and B, P = 0.032), but was markedly reduced by 5 weeks of Mdivi-1 treatment (Fig. 5A and B, P = 0.040). Mitochondrial circumference, an indicator of mitochondrial size, was higher in skeletal muscle from D2-mdx mice compared to WT (Fig. 5C, P = 0.007), but was significantly decreased with Mdivi-1 treatment (Fig. 5C, P = 0.003), indicating Mdivi-1 may have attenuated mitochondrial swelling. Mitochondrial roundness, a parameter indicating fragmented mitochondria, was higher in the D2-mdx model compared to WT (Fig. 5D, P = 0.030), but was significantly attenuated with Mdivi-1 treatment (Fig. 5D, P = 0.017), indicating reduction in mitochondrial fission. Consistently, there was a significantly lower aspect ratio (the ratio of the length of a mitochondrion to its width) in D2-mdx mice compared to the WT group (Fig. 5E, P = 0.035) and 5-week Mdivi-1 treatment had a trend towards significant enhancement of aspect ratio in D2- mdx mice in comparison to the vehicle-treated group (Fig. 5E, P = 0.061), indicating more elongated mitochondria in skeletal muscle of D2-mdx treated with Mdivi-1. Regarding intermyofibrillar mitochondria, there were no statistically significant differences detected among groups with the exception of circumference (Fig. 5F-I). A significant reduction in intermyofibrillar mitochondrial circumference was noted in skeletal muscle with Mdivi-1 treatment (Fig. 5F, P = 0.030).

**Figure 5:**
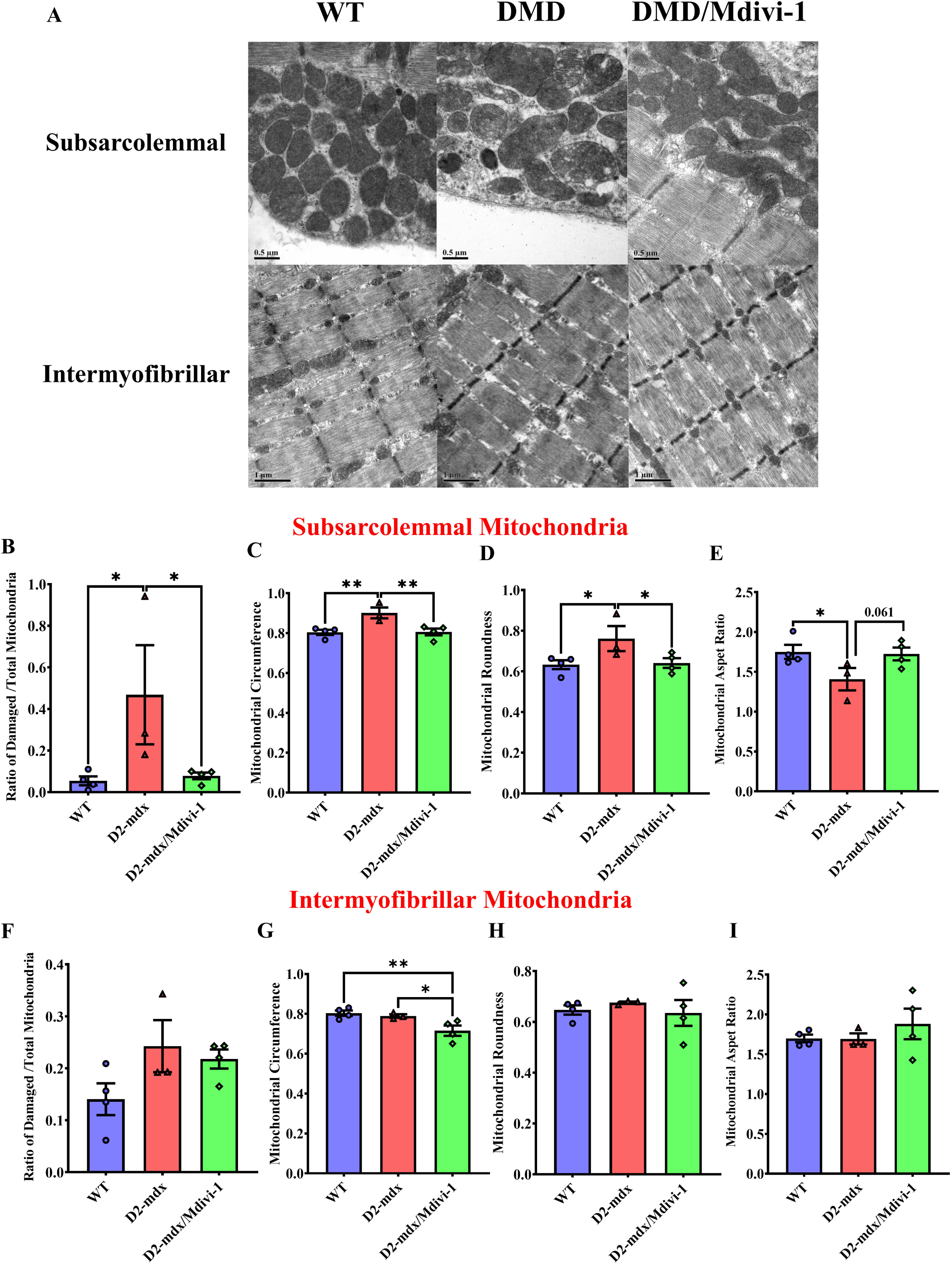
Mdivi-1 treatment improves subsarcolemmal, but not intermyofibrillar mitochondrial morphology in skeletal muscle from D2-mdx mice. A) representative TEM images. B) Ratio of Subsarcolemmal (SS) damaged mitochondria over total mitochondria. C) SS mitochondrial circumference. D) SS mitochondrial roundness. E) SS mitochondrial aspect ratio. F) Intermyofibrillar (IMF) damage mitochondria over total mitochondria. G) IMF mitochondrial circumference. H) IMF mitochondrial roundness. I) IMF mitochondrial aspect ratio. Data are presented as Mean values ± SEM. N= 3-4 mice with 5 TEM images quantified per animal (13,000x magnification). * P < 0.05; ** P < 0.01 significant difference between groups.

### Mdivi-1 treatment improves skeletal muscle mitochondrial respiration in D2-mdx mice

Isolated mitochondria from skeletal muscle of D2-mdx mice exhibited greatly compromised ADP and FCCP-stimulated respiration in comparison to WT mice (Fig. 6A and B, P = 0.009 and 0.030). Although not statistically significant, 5 weeks of Mdivi-1 treatment enhanced ADP and FCCP-stimulated respiration by 93.8% and 92.4% in D2-mdx mice compared to D2-vehicle treated group (Fig. 6A and B, P = 0.061 and 0.171). No difference in basal and state 4 respiration was noted among the three groups. Mitochondrial spare capacity (i.e., maximal respiration rate – basal respiration rate), an important aspect of mitochondrial function, was significantly lower in skeletal muscle from D2-mdx mice than WT mice (Fig. 6C, P = 0.034). However, such difference disappeared in D2-mdx mice after 5 weeks of Mdivi-1 treatment (Fig. 6C). RCR is considered as the single most useful general measure of function in isolated mitochondria. A trend towards significantly lower RCR was found in isolated mitochondrial from skeletal muscle of D2-mdx mice when compared to WT (Fig. 6D, P = 0.095) and, although not significant, RCR was enhanced 80.1% in D2-mdx/Mdivi-1 mice compared to D2-mdx (Fig. 6D).

**Figure 6.**
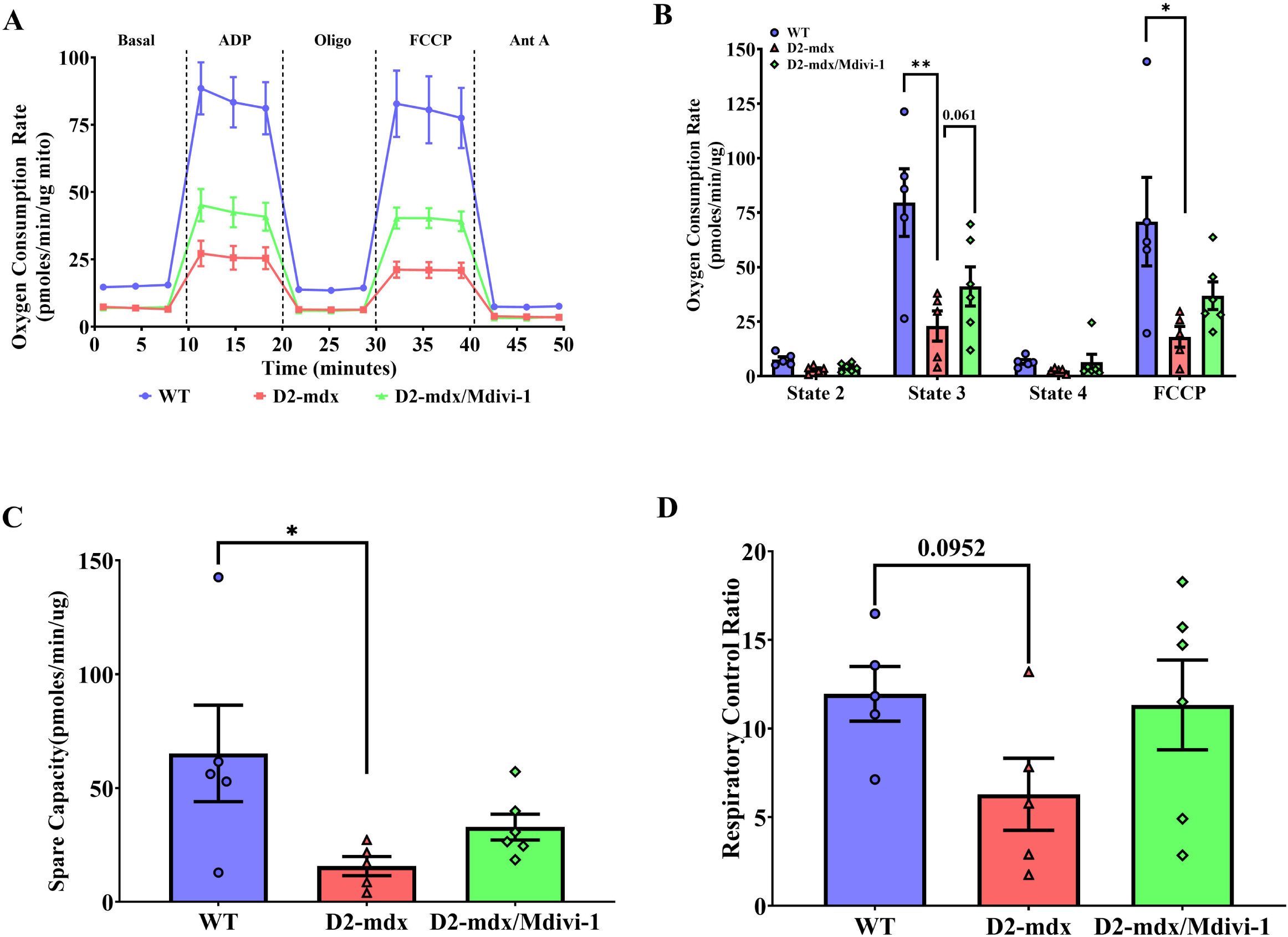
Mdivi-1 treatment has minimal effect on mitochondrial respiration in skeletal muscle from D2-mdx mice. A) Representative graph of oxygen consumption rates (OCR). B) Mitochondrial oxygen consumption rate (OCR). C) spare capacity. D) Respiratory control ratio. Data are presented as Mean ± SEM. N = 5-8 mice per group. * P < 0.05; ** P < 0.01 significant difference between groups.

### Mdivi-1 treatment did not alter mitochondrial H_2_O_2_ production in skeletal muscle from D2- mdx mice

Mitochondria generate H_2_O_2_ during oxidative phosphorylation (74). We next sought to further assess mitochondrial function by measuring mitochondrial H_2_O_2_ production. D2-mdx mice had significantly lower levels of Complex I-, Complex II- and Complex III-supported mitochondrial H_2_O_2_ emission when compared to WT group (Fig. 7A-C, P = 0.014, 0.004 and 0.0001, respectively). However, Mdivi-1 did not significantly alter any of these mitochondrial H_2_O_2_ emission rates in D2-mdx mice (Fig. 7A-C), and Complex II and III- supported mitochondrial H_2_O_2_ emission rates remained significantly lower in D2-mdx/Mdivi-1 mice when compared to WT mice (Fig. 7B-C, P=0.036 and 0.0001, respectively). There were no differences in Pyruvate Dehydrogenase Complex (PDC)-supported mitochondrial H_2_O_2_ emission among the three groups (Fig. 7D).

**Figure 7:**
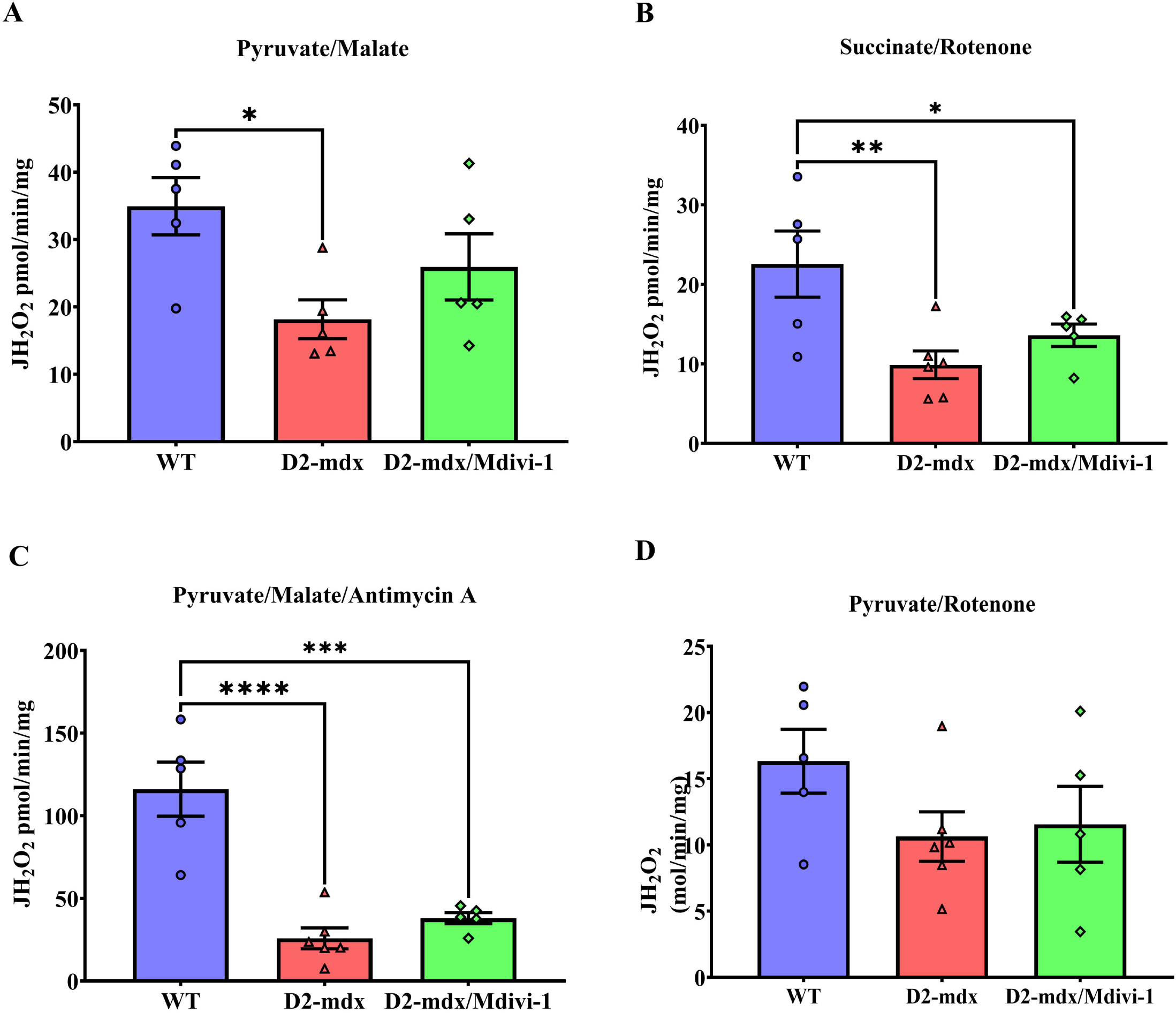
Mdivi-1 treatment did not alter mitochondrial hydrogen peroxide production in skeletal muscle from D2-mdx mice. A) Pyruvate/malate (Complex I-supported mH_2_O_2_ production). B) Succinate/rotenone (Complex II-supported mH_2_O_2_ production). C) pyruvate/malate/antimycin (Complex III-supported mH_2_O_2_ production). and D) pyruvate/rotenone (Pyruvate Dehydrogenase Complex-supported mH_2_O_2_ production). Data are presented as Mean values ± SEM. N= 5-8 mice per group. * P < 0.05; ** P < 0.01; *** P < 0.001; **** P < 0.0001 significant difference between groups.

### Mdivi-1 treatment reduces lipid peroxidation in skeletal muscle from D2-mdx mice

ROS refers to a collection of radical molecules (e.g., hydrogen peroxide (H_2_O_2_), lipid hydroperoxide (LOOH)) (44). Since Mdivi-1 did not improve mitochondrial H_2_O_2_ emission in D2-mdx mice, we next measured other sources of ROS. 4-HNE, a marker for lipid peroxidation (17), was markedly higher in the skeletal muscle from D2-mdx mice than WT counterparts (Fig 8A, P <0.0001), but was significantly reduced with Mdivi-1 treatment (Fig. 8A, P = 0.008). In addition, 4-HNE expression had a significant inverse correlation with grip strength and holding impulse (Fig. 8B and C, r = -0.49 and -0.53, respectively; P=0.016 and 0.012, respectively).

**Figure 8:**
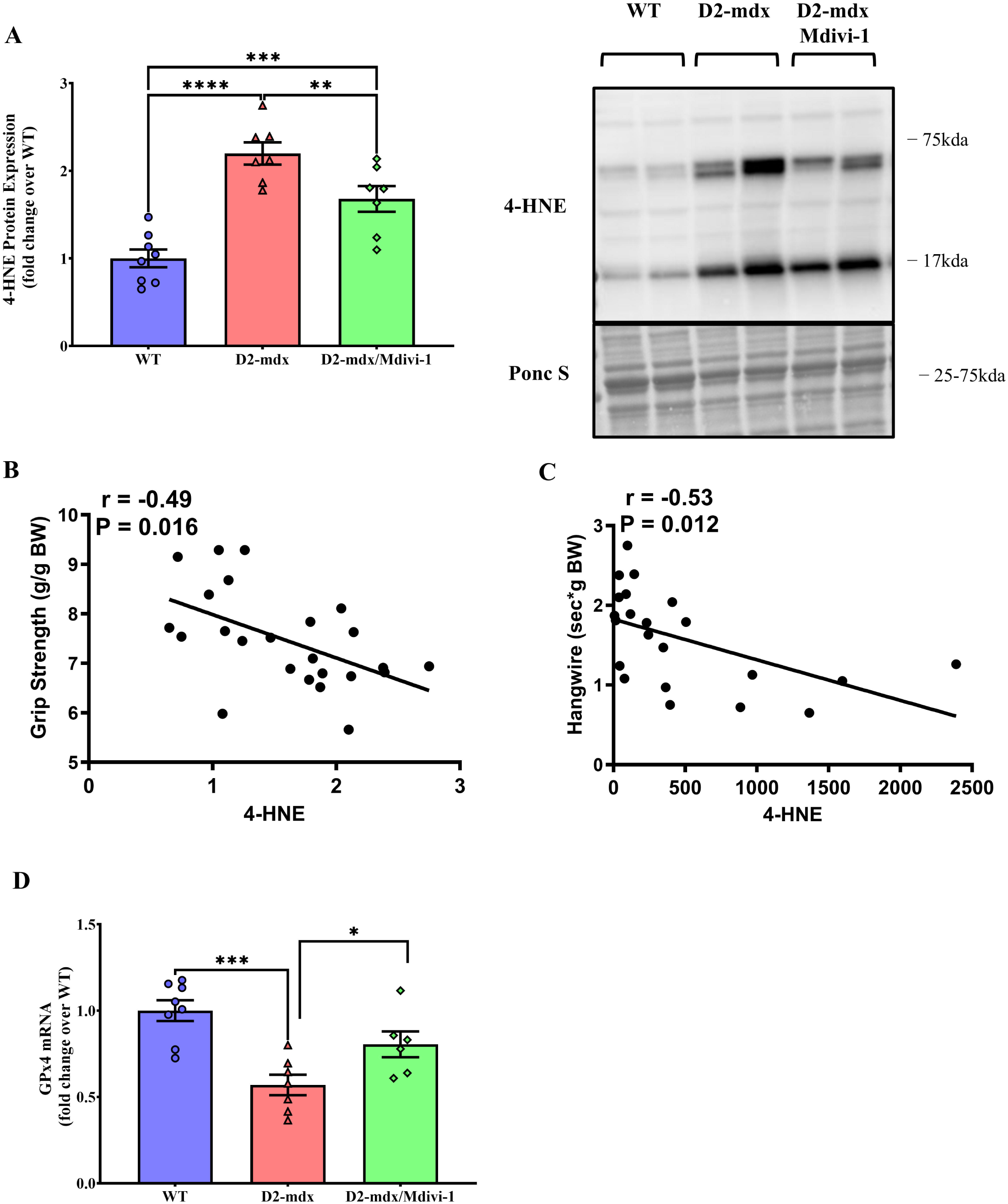
Mdivi-1 treatment reduces lipid peroxidation in skeletal muscle D2-mdx mice. A) 4-HNE protein expression. B) Correlation between 4-HNE protein expression and grip strength. C) Correlation between 4-HNE protein expression and hangwire time. Data are presented as Mean values ± SEM. D) GPx4 mRNA expression. N= 7-8 mice per group. * P < 0.05; ** P < 0.01; *** P < 0.001; **** P < 0.0001 significant difference between groups.

Glutathione peroxidase 4 (GPx4) is an essential antioxidant enzyme that catalyzes the reaction by which LOOH is reduced to its nonreactive hydroxyl metabolite (18, 22). GPx4 mRNA content was lower in skeletal muscle from D2-mdx mice (Fig. 8D, P = 0.0002) but was effectively elevated by Mdivi-1 treatment (Fig. 8D, P = 0.031).

### Mdivi-1 reduces markers of skeletal muscle fibrosis and inflammation D2-mdx mice

Next, we chose to assess the effects of Mdivi-1 on markers of skeletal muscle fibrosis and inflammation. Alpha 1 Type I Collagen (Col1a1) and Fibronectin (FN1) were used as protein markers of fibrosis. There were higher levels of Col1a1 and FN1 protein contents in skeletal muscles from D2-mdx mice when compared to WT controls (Fig. 9A, P = 0.0002 and 0.018, respectively). Mdivi-1 treatment significantly reduced Col1a1 (Fig. 9A, P = 0.001), but not FN1 in D2-mdx mice when compared to the vehicle treated counterparts (Fig. 9A). Furthermore, 4- HNE expression had a significant correlation with Col1a1 protein expression (Fig. 9B, r = 0.55; P=0.008). Lastly, IL-6 mRNA content was higher (Fig. 9C, P = 0.025) in skeletal muscle from D2-mdx mice but was attenuated by Mdivi-1 treatment (Fig. 9C, P = 0.042). In contrast, higher IL-1b mRNA content in skeletal muscle from D2-mdx mice was not affected by Mdivi-1 treatment (Fig. 9D).

**Figure 9:**
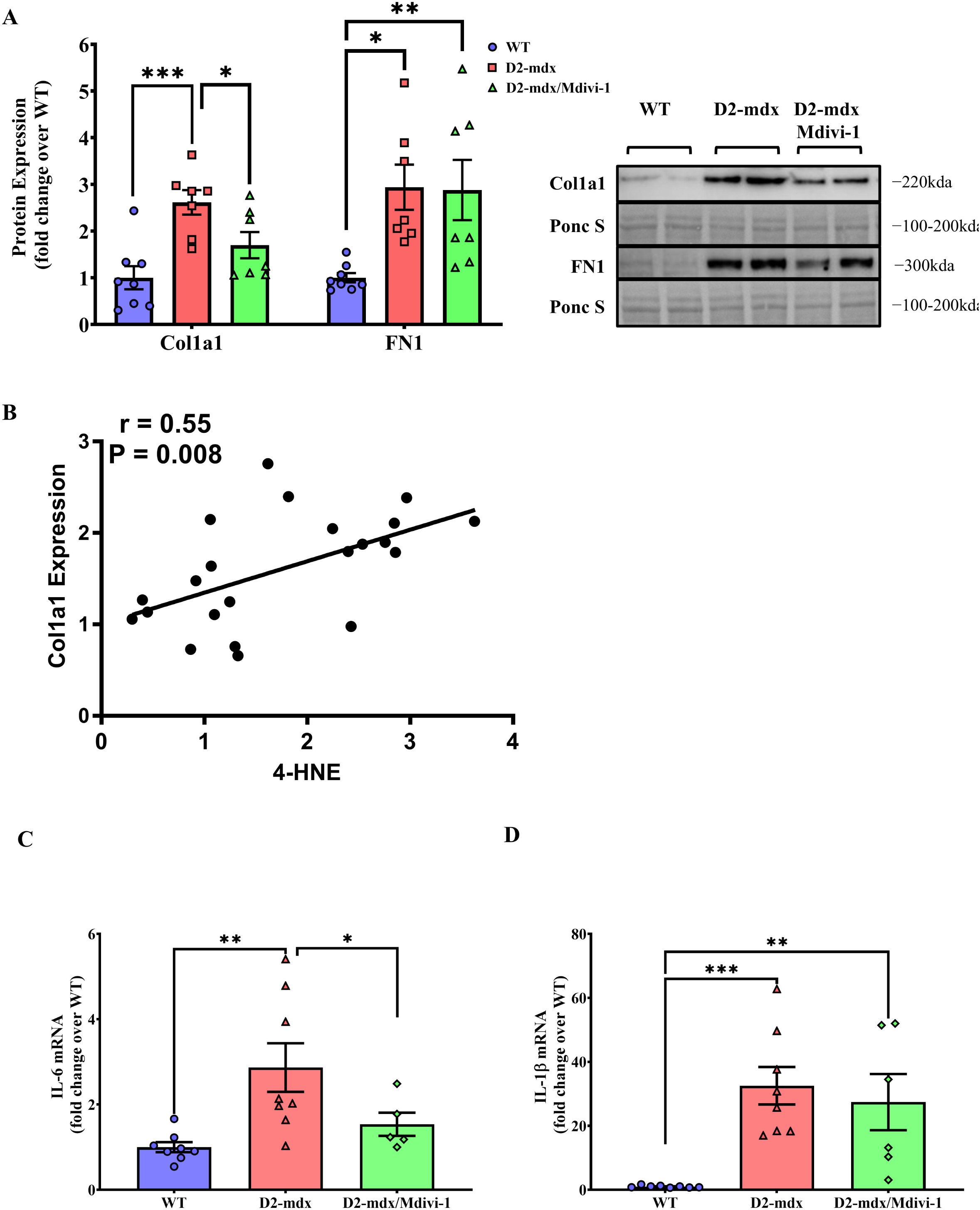
Mdivi-1 treatment improved protein/gene expression of fibrosis and inflammation markers in skeletal muscle from D2-mdx mice. A) Collagen 1 (Col1a1) and Fibronectin 1 (FN1) protein expressions. B) Correlation between 4-HNE protein expression and Col1a1 protein expression. C) IL-6 mRNA expression. D) IL-1β mRNA expression. Data are presented as Mean values ± SEM. N= 6-8 mice per group. * P < 0.05; ** P < 0.01; *** P < 0.001 significant difference between groups.

## DISCUSSION

Despite the recent approval of the first dystrophin gene therapy for DMD, it has only been approved for a subset of patients (those aged 4-5 years old) and at best it can only partly increase dystrophin protein content (31). Given these limitations of the current gene therapy to restore dystrophin, targeting secondary defects (e.g., inflammation, fibrosis) to improve quality of life in DMD patients has been one of the major focuses for guiding therapy development (29). Increasing evidence suggests that mitochondrial dysfunction is an early pathological hallmark of DMD that may culminate in skeletal muscle inflammation, fibrosis, and eventual muscle weakness (33, 50, 51). Here, we demonstrated that Mdivi-1, a pharmacological inhibitor targeting Drp1-mediated mitochondrial fission, was effective in reducing muscle damage, inflammation and fibrosis makers, and improving skeletal muscle strength in D2-mdx mice after 5 weeks of treatment. These improvements are associated with improved mitochondrial morphology and reduced lipid peroxidation. To our knowledge, this study is the first to investigate Drp1-mediated mitochondrial fission, a key process in maintaining mitochondrial quality and function, as a potential drug target for developing novel therapies to alleviate myopathy in DMD.

The main finding of the present study is that D2-mdx mice treated with Mdivi-1, a pharmacological inhibitor of Drp1, showed significantly improved muscle strength and reduced muscle damage compared to their vehicle-treated counterparts. Consistent with our findings, Rexius-Hall et al. recently reported that Mdivi-1 increased contractility generated by engineered muscle fibers *in vitro* (59). In DMD, muscle weakness and dysfunction are largely due to the presence of elevated fibrosis and chronic inflammation within the skeletal muscle tissue, which results from the degeneration of muscle fibers caused by the lack of dystrophin protein (35). Our findings of reduced expressions of Col1a1 and IL-6 in skeletal muscle tissues from Mdivi-1 treated DMD mice suggest that the improved skeletal muscle strength observed in D2-mdx mice may be due to the attenuated fibrosis and inflammation. The extracellular matrix (ECM) is primarily comprised of collagens (>80%) with the most common type of collagen in skeletal muscle tissue as Collagen I (36). Skeletal muscle from DMD models normally exhibit high levels of muscle collagen content, leading to fibrosis, compromised muscle quality and strength (49). It has been shown that targeting fibrosis in DMD models was effective to improve muscle strength and function (16, 32, 68). In addition, we observed reduced IL-6 gene expression in skeletal muscle tissue from D2-mdx mice treated with Mdivi-1. IL-6 is an inflammatory cytokine that is chronically elevated in DMD, which can promote inflammation and necrosis, leading to fibrosis (53). In line with this, studies using anti-IL-6 receptor antibody in mdx mice have shown attenuated muscle fibrosis, atrophy and improved muscle regeneration and strength (52, 70). Overall, our data suggest that Mdivi-1 treatment may reduce IL-6 production in skeletal muscle, which subsequently attenuate inflammation and fibrosis, leading to improved muscle strength in D2-mdx mice. It is worth noting that we did not perform direct evaluations of fibrosis with histology (e.g. hematoxylin and eosin (H&E) and Masson’s trichrome staining) in this study, which is one of the limitations. Future studies should warrant such assessment of fibrosis to provide definitive evidence as to whether Mdivi-1 can effectively reduce fibrosis in skeletal muscle from DMD mice.

In the present study, several regulatory markers of mitochondrial fission, including Drp1(Ser616) phosphorylation, Drp1 and Fis1 protein content, were significantly higher in skeletal muscle from D2-mdx mice. These finding are in agreement with previous studies using various models of DMD, demonstrating that skeletal muscle mitochondrial dynamics was imbalanced in a manner that shifted towards mitochondrial fission with excessive activation of Drp1-mediated fission machinery at young age (9-11 weeks) (30, 50, 51, 61, 62). Importantly, 5 weeks of Mdivi-1 administration was successful in reducing Drp1-mediated mitochondrial fission in D2-mdx mice, suggesting inhibited Drp1 activity by Mdivi-1 likely contributed to the improvements in skeletal muscle in D2-mdx mice. Since discovered by Cassidy-Stone and colleagues in 2008 (14), Mdivi-1 is by far the most accessible and frequently studied pharmacological inhibitor of primary mitochondrial fission protein Drp1 (45, 63), with the evidence that Mdivi-1 inhibits mitochondrial fission, leading to elongated mitochondria in various types of cells. More importantly, the therapeutic potential of Mdivi-1 has been extensively reported in various neurodegenerative disease models such as Amyotrophic Lateral Sclerosis and Alzheimer’s disease (46, 60), highlighting its clinical potential (45). However, two recent studies found no evidence that Mdivi-1 acts as a mitochondrial fission inhibitor and identified off-target effects (7, 37). The discrepancy in results from different studies may be attributed to different protocols of Mdivi-1 treatment (e.g., concentration, duration and cell lines). Our Drp1 phosphorylation data, coupled with mitochondrial morphology data from TEM images, demonstrated that Mdivi-1 is an effective inhibitor of Drp1 and mitochondrial fission in skeletal muscle cells. Our findings are consistent with previous studies utilized muscle cell line in vitro and/or skeletal muscle tissues in vivo (40, 46, 59), suggesting Mdivi-1 is an effective inhibitor targeting Drp1-mediated mitochondrial fission in skeletal muscle.

Furthermore, we found a significantly lower ratio of LC3B II/I in skeletal muscle from D2-mdx mice, suggesting that autophagic flux was blunted in dystrophin-deficient muscles. This finding agrees with several studies using skeletal muscle samples from mdx mice and DMD patients (6, 19, 39, 65), but is contradictory to a recent study (50). Moore et al., reported no changes in autophagic protein markers in mdx mice in comparison to the age-matched WT controls (50). The discrepancy in the findings of LC3B may be due to the utilization of different mouse models; Moore and colleagues used the B10.mdx mouse model at young age, which presents mild myopathy, whereas we utilized the more severe D2-mdx model. In fact, Spitali et al. reported lower LC3B II/I ratio in skeletal muscle from 16-week-old mdx mice, a stage with more severe DMD symptoms (65). In addition, the discrepancy may also arise from the use of different muscles for detecting LC3B protein. Moore et al. used quadriceps for immunoblotting, whereas we analyzed protein expression alterations in gastrocnemius. Unexpectedly, Mdivi-1 treatment improved LC3B II/I ratio in mdx mice, indicating an enhancement in autophagic flux to allow irreversibly damaged cellular components to be cleared rather than accumulating. Future studies are needed to validate the effects of Drp1-mediated mitochondrial fission inhibition on autophagic flux and, if validated, to explore the mechanism by which Drp1-inhibition is involved in the promotion of autophagic flux.

The balance of mitochondrial dynamics controls mitochondrial morphology. Compared to WT, subsarcolemmal (SS) mitochondria exhibited more damage and fragmented shapes (e.g., elevated roundness and reduced aspect ratio) than intermyofibrillar (IMF) mitochondrial in the skeletal muscle from D2-mdx mice. This is consistent with previous studies demonstrating that SS mitochondria are more sensitive and prone to damage due to muscle disuse and atrophy (38). In the current study, the morphology of SS mitochondria in mdx mice was significantly improved by Mdivi-1 treatment with a notable shift towards elongated mitochondria. These findings corroborate with the alterations in proteins responsible for mitochondrial fission, validating Mdivi-1 indeed inhibited Drp1-mediated mitochondrial fission. Interestingly, our study found that 5 weeks of Mdivi-1 treatment did not improve IMF mitochondrial morphology. We speculate that insufficient amount of Mdivi-1 may have reached to IMF mitochondria due to relatively deeper location within myofibers compared to SS mitochondria.

ROS accumulation is a common feature in skeletal muscle from mdx mice due to the compromised oxidative phosphorylation in mitochondria and plays an important role in the pathogenesis of DMD (33, 64). Excessive ROS accumulation is tightly linked to inflammation, fibrosis, and necrosis in skeletal muscle from DMD. There are several sources of ROS, including hydrogen peroxide (H_2_O_2_), hydroxyl radical (OH.), and lipid hydroperoxide (LOOH). Hughes et al., found mitochondria-derived H_2_O_2_ emission during oxidative phosphorylation was higher in skeletal muscle from 4-week-old D2-mdx mice compared to the WT controls (33). Surprisingly, we found reduced mH_2_O_2_ emission in mdx mice and there was no significant effect of Mdivi-1 on mH_2_O_2_ emission regardless of substrates. Our finding is consistent with a previous study, in which lower mH_2_O_2_ production was reported in skeletal muscle from 6-week-old male mdx mice compared to WT (27). These contradictory findings regarding mH_2_O_2_ emission may be due to the different age of mdx mice used in the studies. Hughes et al. used 4-week-old mdx mice, which may have developed early mitochondrial dysfunction with compromised mitochondrial respiration capacity and elevated mH_2_O_2_ emission. Subsequently, a compensatory adaptation may occur in dystrophin-deficient skeletal muscle at later stage (6-9-week-old) to counteract oxidative stress with enhanced mH_2_O_2_ scavenging capacity (e.g., antioxidant system) in response to oxidative stress (Godin et al., 2012).

In contrast to mH_2_O_2_ emission, 4-HNE, a marker of lipid peroxidation, was higher in D2- mdx mice compared to WT. This is consistent with multiple studies finding markers of lipid peroxidation elevated in plasma and skeletal muscle biopsies from DMD patients (20, 28, 34, 47). In addition, it has also been shown that increased production of LOOH (5, 22, 54), a byproduct of lipid peroxidation, but not H_2_O_2_ (23) was associated muscle wasting and weakness. More importantly, our study found that Mdivi-1 treatment effectively reduced lipid peroxidation in skeletal muscle from 9-week-old D2-mdx mice. While no study has been done in targeting lipid peroxidation in DMD, recent studies reported that inhibition of LOOH protected muscle loss and improved muscle strength, supporting that LOOH plays an important role in regulating skeletal muscle mass and function (22, 54). It is also possible that mH_2_O_2_ emission may be elevated at the early age in D2-mdx model, which may initiate the cascade of other ROS generations (e.g., LOOH) at the later age. Regardless, a future time-course study should be conducted to identify different sources of ROS, including H_2_O_2_ and LOOH in skeletal muscle from different ages of mdx mice, to determine the optimal time window for targeting these different ROS sources for developing additional therapies to treat myopathy.

Our study has some limitations. First, our study only investigated one dose (40mg/kg BW) of Mdivi-1 administration. Future studies should consider assessing the efficacy of Mdivi-1 using different doses and longer durations to validate its potential therapeutic efficacy and safety. In fact, we found partial restoration of several markers in mitochondrial and skeletal muscle health (e.g., respiration, CK, muscle strength). We anticipate that there may be more robust improvements with longer duration of Mdivi-1 treatment. Second, although grip strength and hang wire impulse tests are commonly utilized for *in vivo* assessment of muscle strength in mice, they are both indirect measures. Future studies should utilize a more direct assessment of skeletal muscle strength, such as *ex vivo* contractile assessments of isometric force. Finally, Mdivi-1 was delivered systemically via intraperitoneal injection. Therefore, it is unclear if the improvements on grip strength and muscle damage seen in our study can be directly attributed to the enhancement in skeletal muscle mitochondria. Future studies may consider attempting intramuscular injection of Mdivi-1 in the D2-mdx mouse model of DMD.

In conclusion, we provide evidence that inhibition of Drp1-mediated mitochondrial fission *in vivo* using Mdivi-1 effectively enhanced muscle strength and mitigated overall muscle damage in D2-mdx mouse model of DMD. We further demonstrated that these improvements are associated with, and may partly be due to, improved skeletal muscle mitochondrial integrity, leading to attenuated lipid peroxidation and subsequently reduced inflammation and fibrosis. Together, these results add significant knowledge to the growing research field regarding the contribution of mitochondria, a critical but underappreciated organelle, to the pathophysiology of DMD and identify a novel target for the potential therapeutics of targeting Drp1-mediated mitochondrial fission to improve quality of life in DMD patients.

## ACKNOWLEDGMENTS

This study was supported by grants from the National Institutes of Health (R15DK131512, K.Z.).

This publication was made possible by the Electron Microscopy Core Facility at the UMass Chan Medical School through grant S10 OD025113-01 from National Institutes of Health. We would like to thank Drs. Changmeng Cai and Alexey Veraksa for their support and feedback. We would also like to thank Dr. Gregory Hendricks and Mr. Keith Reddig for their technical support.

