## Supplemental Tables for "Inhibition of Mitochondrial Fission Protein Drp1 Ameliorates Myopathy in the D2-mdx Model of Duchenne Muscular Dystrophy"

**Supplemental Table 1**: Complete list of antibodies

| Antibody | Company | Dilution | Catalog # |
| --- | --- | --- | --- |
| Fis1 | Abnova | 1:1000 | H00051024-M01 |
| Total Drp1 | Cell Signaling | 1:1000 | 8570S |
| P-Drp1(ser616) | Cell Signaling | 1:1000 | 3455S |
| MFF | ProteinTech | 1:1000 | 17090-1-AP |
| MTP18 | ProteinTech | 1:1000 | 14257-1-AP |
| OPA1 | Cell Signaling | 1:1000 | 80471S |
| Mfn1 | Abcam | 1:1000 | AB104274 |
| Mfn2 | Abcam | 1:1000 | AB56889 |
| OXPHOS Rodent Cocktail | Abcam | 1:5000 | AB110413 |
| Bnip3 | Cell Signaling | 1:1000 | 3769S |
| LC3B | Cell Signaling | 1:1000 | 4108S |
| Parkin | Cell Signaling | 1:1000 | 32833S |
| PGC1α | Novus Biological | 1:1000 | NBP104676 |
| P62 | Cell Signaling | 1:1000 | 5114S |
| Pink1 | Santa Cruz | 1:500 | 38CT20.8.5 |
| Col1α1 | Cell Signaling | 1:1000 | 72026S |
| FN1 | Cell Signaling | 1:1000 | 63779S |
| VDAC | Cell Signaling | 1:1000 | 4866S |
| HRP-linked anti-rabbit | Cell Signaling | 1:5000 | 7074P2 |
| HRP-linked anti-mouse | cell Signaling | 1:5000 | 7076P2 |

**Supplemental Table 2:** Taqman probes used for the real-time quantitative polymerase chain reaction

| **Gene Name** | **Assay Identifier** |
| --- | --- |
| *IL-6* | Mm00446190_m1 |
| *IL-1β* | Mm00434228_m1 |
| *GAPDH* | Mm99999915_g1 |

Abbreviations: IL-6, Interleukin 6; IL-1β, Interleukin 1 beta; GAPDH, glyceraldehyde-3 phosphate dehydrogenase.

**Supplemental Table 3:** Primers used for the real-time quantitative polymerase chain reaction

| **Gene Name** | **Primers** |
| --- | --- |
| *GPx4* | F: TCTGTGTAAATGGGGACGATGC  R: TCTCTATCACCTGGGGCTCCTC |
| *β-Actin* | F: GACGGCCAAGTCATCACTATTG  R: CCACAGGATTCCATACCCAAGA |

Abbreviations: GPx4, Glutathione peroxidase 4.
